# Growth Cone-Localized Microtubule Organizing Center Establishes Microtubule Orientation in Dendrites

**DOI:** 10.1101/841759

**Authors:** Xing Liang, Marcela Kokes, Richard Fetter, Melissa A. Pickett, Maria D. Sallee, Adrian W. Moore, Jessica Feldman, Kang Shen

## Abstract

A polarized arrangement of neuronal microtubule arrays is the foundation of membrane trafficking and subcellular compartmentalization. Conserved among both invertebrates and vertebrates, axons contain exclusively “plus-end-out” microtubules while dendrites contain a high percentage of “minus-end-out” microtubules, the origins of which have been a mystery. Here we show that the dendritic growth cone contains a non-centrosomal microtubule organizing center (ncMTOC), which generates minus-end-out microtubules along outgrowing dendrites and plus-end-out microtubules in the growth cone. RAB-11-positive recycling endosomes accumulate in this region and are responsible for localizing the microtubule nucleation complex γ-TuRC. The MTOC tracks the extending growth cone by kinesin-1/UNC-116-mediated endosome movements on distal plus-end-out microtubules and dynein-mediated endosome clustering near MTOC. Critically, perturbation of the function or localization of the MTOC causes reversed microtubule polarity in dendrites. These findings unveil the dendritic MTOC as a critical organelle for establishing axon-dendrite polarity.

---

Microtubules (MTs) are nucleated and organized by microtubule organizing centers (MTOCs). These distinct cellular sites nucleate, anchor, and stabilize MT minus ends, thereby creating cell-type specific MT organization and protecting MT minus ends against depolymerization. However, mature neurons lack a MTOC and MT arrays are organized with a stereotypical plus-end-out (PEO) orientation in axons, while dendrites contain predominantly minus-end-out (MEO) or both MEO and PEO MTs in invertebrate and vertebrate neurons, respectively^1–3^. While it is perhaps easier to imagine how PEO MTs might arise as an extension of the plus-end-cortical MT arrangement during cell division, the origin of MEO MTs is harder to grasp. Several mechanisms have been proposed to create a bias towards MEO MTs found in dendrites, including microtubule sliding^4,5^, minus end growth^6^, and Golgi outpost-mediated MT nucleation and organization^7^. However, direct evidence for how MEO MTs are first established in the dendrite has been lacking^8^.

To investigate the establishment of MT organization during dendrite outgrowth, we visualized the endogenous localization of the MT plus-end tracking protein EBP-2::GFP/EB1 in developing PVD dendrites in *C. elegans*. PVD is a sensory neuron with stereotypical morphology of its axon and non-ciliated dendrites^9^. We find that early PVD morphogenesis is temporally and spatially stereotyped: the axon always grows out first, followed by the anterior and then posterior dendrite, with all emerging neurites oriented in the direction of their mature neurite pattern (Extended Data Fig. 1a). We previously showed that the mature anterior dendrite contains largely MEO MTs while the posterior dendrite and axon have PEO MTs^10^. During the outgrowth of the anterior dendrite, we found that numerous EBP-2 comets emerged from a single region within the distal neurite, suggesting the presence of a dendritic growth cone MTOC, or dgMTOC (Fig.1a, Extended Data Movie 1). To characterize the directionality of these EBP-2 comets, we plotted the frequency of PEO and MEO MTs at interval distances in the anterior dendrite, setting the center of the apparent dgMTOC as zero (Fig. 1b). Near the most distal region of the dendrite (∼3-6 µm from the dendrite tip), almost all the EBP-2 comets move towards the distal tip indicating PEO microtubules, while nearly all the comets along the shaft of the anterior dendrite move towards the cell body thus highlighting only MEO MTs in this region (Fig. 1a, b). The emergence of a high frequency of EBP-2 comets from a discrete region near the dendrite tip (mean=0.44 comets/s ±0.11 (SD) in a 3.2 µm region), which also encompasses a transition zone of MT directionality, is consistent with the presence of a local MTOC (Fig. 1a, b). This MT organization is apparent as soon as a morphologically distinct anterior dendrite has emerged from the cell body and remains throughout the outgrowth of the anterior dendrite (data not shown). In addition, the number of comets generated near the dendrite tip greatly exceeds comets from any other discrete point or region in the dendrite or from the cell body throughout the morphogenesis of the anterior dendrite, indicating that the dgMTOC is the only apparent MTOC. The axon and posterior dendrite, which contain predominantly PEO MTs, do not contain similar structures (Fig. 1a and Extended Data Fig. 1b). Significantly, we find evidence for a similar dgMTOC near the tip of outgrowing *Drosophila melanogaster* Class I sensory vpda dendrites (Extended Data Fig. 1c, Movie 2), suggesting a conserved mechanism for the genesis of MEO dendritic MTs.

**Fig. 1.**
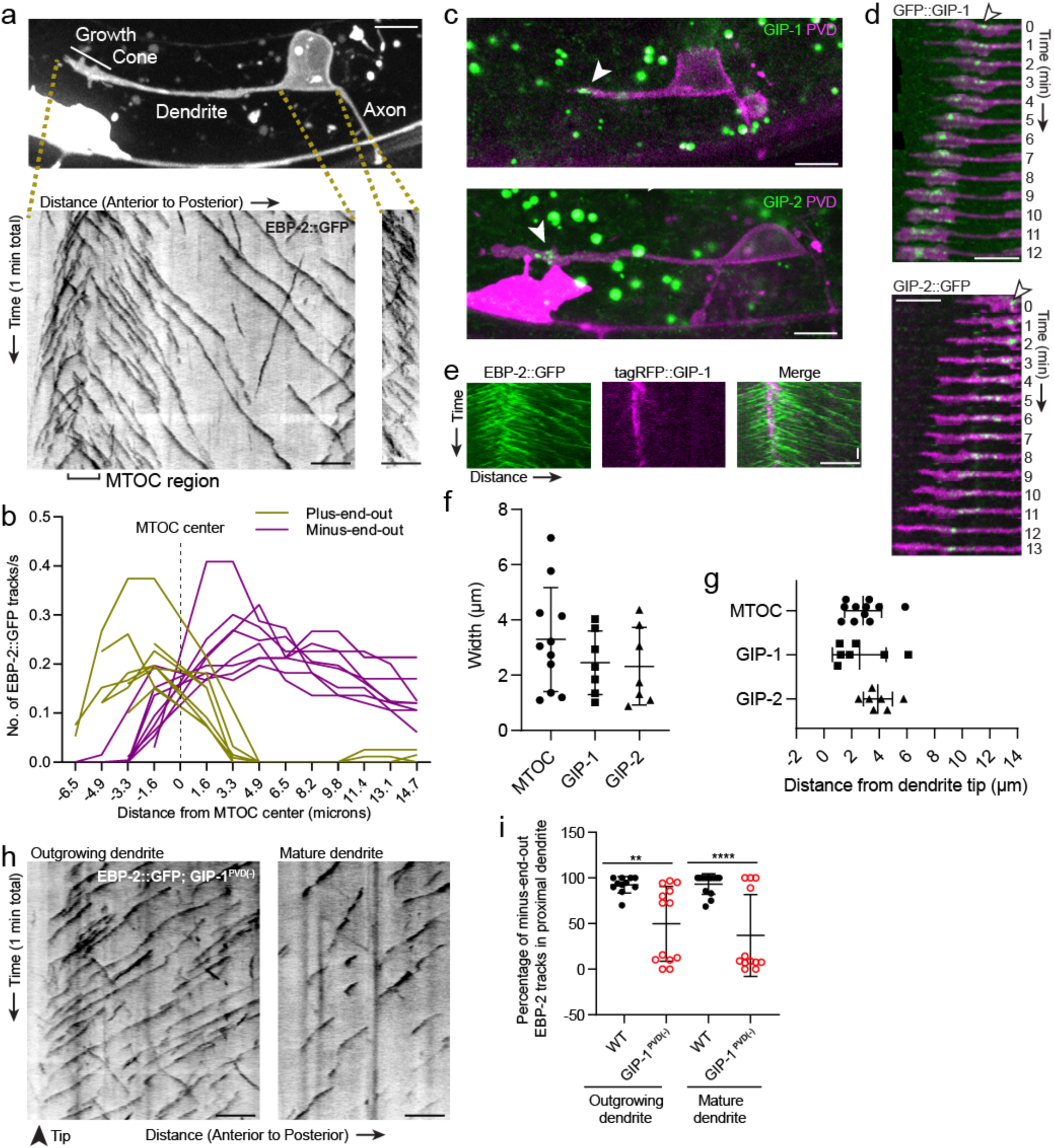
An active MTOC localizes to the growth cone of the outgrowing PVD dendrite. (**a**) Top: A labeled example of PVD morphology during early anterior dendrite outgrowth. Bottom: Kymographs of EBP-2::GFP in an outgrowing dendrite (left) and axon (right). (**b**) Number of plus-end-out and minus-end-out EBP-2 tracks per second at interval distances from the MTOC center (dashed line) in 8 individual animals. (**c**) Endogenously-tagged GFP::GIP-1 (top) and GIP-2::GFP (bottom) localization in an outgrowing PVD dendrite. White arrowhead, GIP cluster Unrelated structures outside of PVD include: gut granules (large green spots), HSN cell body (bright magenta region) (**d**) GIP-1 (top) and GIP-2 (bottom) localization at different time points during live imaging. White arrowhead, GIP cluster. (**e**) Kymograph of EBP-2::GFP and tagRFP::GIP-1 in the growth cone region, horizontal scale bar, 10s (**f-g**) Quantification of the width (f) and shortest distance from the GIP or MTOC region to the dendrite tip (g). MTOC region was identified using EBP-2-GFP kymographs as in A. Error bars represent the standard deviation (SD). (**h**) Kymograph of EBP-2::GFP following PVD-specific depletion of GIP-1 in an outgrowing (left) and mature (right) dendrite. (**i**) Quantification of MT polarity in the proximal dendrite following PVD-specific GIP-1 depletion. Scale bar, 5 µm. All images are lateral views oriented with anterior to the left and ventral down.

Microtubules are nucleated by the conserved γ-tubulin ring complex (γ-TuRC), a ring of γ-tubulin complex proteins (GCPs) that templates the assembly of new MTs^11,12^. γ-TuRCs localize to MTOCs, including the centrosome, the best studied MTOC which generates MTs from within its pericentriolar material (PCM) in dividing animal cells to form the mitotic spindle^13^. To understand the molecular components of the dgMTOC, we examined the localization of endogenous GFP::GIP-1/GCP3 and GIP-2::GFP/GCP2 – core components of the γ-TuRC that mark MT minus ends – during dendrite outgrowth. We found that GIP-1 and GIP-2 consistently localized near the dendrite tip in a cluster of punctate structures which track with the growth cone in the outgrowing anterior dendrite (Fig. 1c, d). GIP-1 precisely localized to the site from which EBP-2 comets originated (Fig. 1e) and quantification of the distribution and location of GIP-1 and GIP-2 clusters showed that their location closely match the dgMTOC regions defined by EBP-2 comets (Fig. 1f, g). These γ-TuRC clusters are specific to the outgrowing anterior dendrite since no GIP-1 or GIP-2 clusters were found in the cell body, axon, or posterior dendrite during their outgrowth (Fig. 1c, Extended Data Fig. 1d, e), consistent with the lack of an apparent MTOC in these structures. Furthermore, growth cone-localized GIP-2 was visible as soon as the anterior dendrite emerged (Extended Data Fig. 1f) but absent near the tip of mature dendrites (Extended Data Fig. 1g), suggesting that the dgMTOC plays a role in the establishment of MT organization during dendrite development. Significantly, GIP-2 also localized close to the dendrite tip during dendrite outgrowth in the *C. elegans* DA9 motor neuron (Extended Data Fig. 1h), further supporting a conserved role for the dgMTOC in establishing dendritic MEO MTs.

To investigate the significance of the dgMTOC and directly test if γ-TuRC is required for the dgMTOC activity, we performed conditional knockdown of GIP-1 in the PVD lineage using the ZIF-1/ZF degradation system previously established in *C. elegans*^14^. We inserted a ZF tag into the endogenous *gip-1* locus, enabling the controlled degradation of endogenous GIP-1 upon expression of the E3 ubiquitin ligase substrate-recognition subunit ZIF-1. Expression of ZIF-1 in the developing PVD lineage led to a dramatic reduction of punctate GFP::GIP-1 signal in ∼80% of outgrowing PVD anterior dendrites (GIP-1^PVD(-)^, Extended Data Fig. 1i). We then examined EBP-2 comets in GIP-1^PVD(-)^ animals, which revealed a striking reversal of anterior dendrite MT polarity leading to PEO orientation in both outgrowing and mature dendrites in about half of the animals (Fig. 1h, i). Unlike in wild type (wt) animals where the majority of EBP-2 comets were generated by the dgMTOC, in GIP-1^PVD(-)^ animals, many dendritic comets originated from the cell body (Fig. 1h, Extended Data Movie 3). The incomplete penetrance of this phenotype in GIP-1^PVD(-)^ animals is likely due to partial knockdown of GIP-1, as shown by the bimodal distribution of the MT polarity phenotype (Fig. 1i). These results indicate that γ-TuRC is a critical component of the dgMTOC and that the dgMTOC is essential for establishing MEO MTs in both developing and mature dendrites.

To determine the subcellular ultrastructure of the dgMTOC, we performed serial sectioning electron microscopy (EM) reconstruction of developing distal PVD dendrites. We began by reconstructing MTs to identify the dgMTOC region in our sections. In our EM series, numerous short, staggered MTs were observed with their ends unambiguously identified. While EBP-2 labels dynamic MTs, EM reconstruction reveals total MTs. Consistent with our EBP-2 results, the highest MT density was found ∼2-4 um from the dendritic tip, consistent with the presence of an MTOC in this region (Fig. 2a-c). To further confirm the location of the MTOC, we performed time-lapse imaging on worms expressing GFP::TBA-1/α-tubulin in PVD, which allowed us to track MT shrinkage and pausing in addition to growth events. GFP::TBA-1 was enriched in a region immediately behind the growth cone (Extended Data Fig. 2a, b), suggesting higher numbers of MTs in this region. The relatively low number of MTs in *C. elegans* neurons^15^ allowed us to identify individual MTs on kymographs. This analysis also revealed an apparent dgMTOC from which presumed PEO MTs extended distally and MEO MTs extended proximally (blue or red lines, respectively, in Fig. 2d). Consistent with the direction of EBP-2 comets, we observed numerous polymerization events originating from the dgMTOC and directed towards the cell body (red lines in Fig. 2d). A similar frequency of depolymerization events was also observed in this region (green lines in Fig. 2d, Fig. 2e). Notably, 80±4.5% (SEM, n=7) of all polymerization events were immediately followed by depolymerization events (rather than a pause) in this region (Fig. 2f), indicating that the majority of dgMTOC-derived MEO MTs are highly dynamic.

**Fig. 2.**
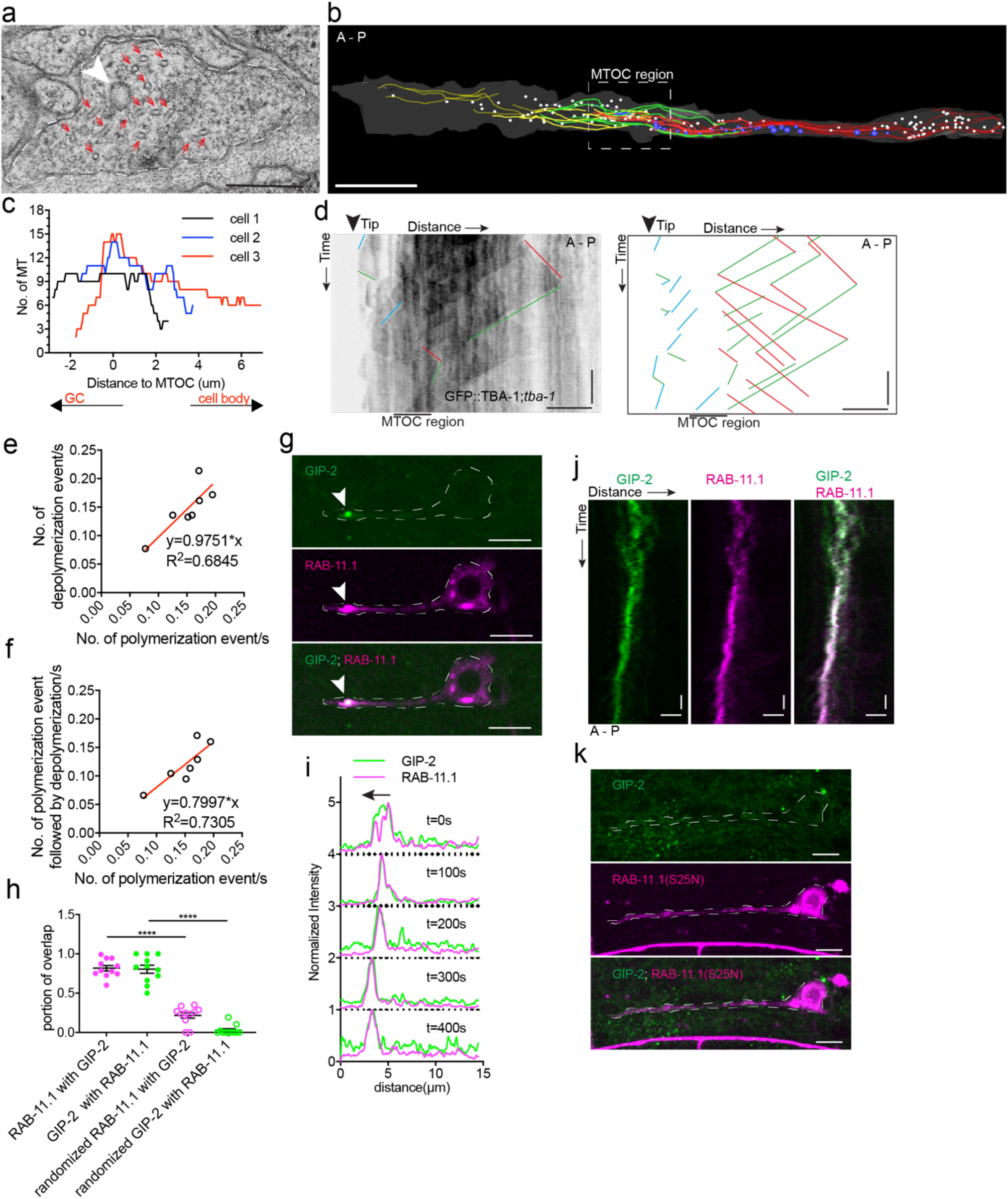
γ-TuRC localizes to RAB-11.1 endosomes to form the dgMTOC. (**a**) Electron micrograph of the dendritic growth cone. Red arrows, MTs; white arrowhead, clear-core vesicle; scale bar, 0.2 μm. (**b**) Reconstruction of serial electron microscopy sections through the anterior dendrite. Yellow lines, MTs in the distal dendrite; green lines, MTs crossing the predicted MTOC region; red lines, MTs in the proximal region; white circles, clear-core vesicles; blue circles, dense-core vesicles; scale bar, 1 μm. (**c**) MT number distribution determined by serial electron microscopy in the anterior dendrite growth cone region in three PVD neurons during development. (**d**) Kymograph of GFP::TBA-1 in the growth cone region. The GFP::TBA-1 was expressed in a *tba-1* null mutant background to get a better incorporation of the GFP::TBA-1. Blue lines, growing PEO MTs in distal region; red lines, growing MEO MTs in proximal region; green lines, retracting MTs; horizontal scale bar, 5 μm; vertical scale bar, 10 s. (**e**) Frequency of polymerization relative to depolymerization events in the proximal region of the anterior dendrite (n=7). (**f**) Frequency of polymerization events followed by depolymerization events relative to total polymerization events in the proximal region of the anterior dendrite (n=7). (**g**) GIP-2 and RAB-11.1 colocalization in the dendritic growth cone region. Scale bar, 5 μm. (**h**) Quantification of GIP-2 and RAB-11.1 fluorescence overlap in the growth cone region. (**i**) Normalized intensity of GIP-2 and RAB-11.1 from the dendritic tip along the dendrite shaft to the cell body at different time points. Black arrow, direction of GIP-2 and RAB-11.1 movement. (**j**) Kymograph of GIP-2::GFP and mCherry::RAB-11.1 in the growth cone region. Horizontal scale bar, 2 μm; vertical scale bar, 10 s. (**k**) GIP-2 localization in worms overexpressing RAB-11.1(S25N) dominant negative mutant. Scale bar, 5μm. A, anterior; P, posterior. Images in g and k were taken in the *glo-1*(*zu391*) mutant background to reduce the gut granule signal.

Serial EM reconstruction showed no evidence of a centrosome or Golgi outpost in the dgMTOC area. Instead, numerous clear- and dense-core vesicles were found in between the MT arrays (Fig. 2a, b). In particular, clusters of clear-core vesicles (Fig. 2b, white) were found in the region with the highest MT numbers suggesting that GIP-1 and GIP-2 might localize to these vesicles to form the dgMTOC. To investigate the molecular identity of these vesicles, we expressed GFP-tagged markers to label different vesicles and membrane compartments. Synaptic vesicles are the most abundant clear-core vesicles in neurons, however, the synaptic vesicle marker RAB-3 which robustly localized to the axon^16^, was not enriched in the dendritic growth cone region (Extended Data Fig. 2c). Consistent with our EM results, the early/medial Golgi cisternae markers RER-1^17^ and AMAN-2/alpha-mannosidase 2, the latter of which localizes to Golgi outpost MTOCs at mature dendrite branch sites in *D. melanogaster*^7^ were only present in the cell body but not in the growth cone (Extended Data Fig. 2d, e). Surprisingly, the late-Golgi compartment marker RAB-6.2^18^ was enriched in the growth cone in a localization pattern similar to that of GIP-1 and GIP-2 (Extended Data Fig. 2d and e). In addition to localizing to Golgi stacks, RAB-6.2 also plays a role in recycling cargo molecules from endosomes to the trans-Golgi network in neurons^18^. Therefore, we further examined recycling endosomes and found that the recycling endosome marker RAB-11.1 co-localized and moved together with RAB-6.2-labeled vesicles in the growth cone (Extended Data Fig. 2f, g). Interestingly, RAB-11.1 also showed striking co-localization with GIP-2 (Fig. 2g), with 81.7±3.5% (SEM, n=11) of RAB-11.1 fluorescence overlapping with GIP-2 and 80.6±5.2% (SEM, n=11) reciprocal GIP-2 fluorescence overlapping with RAB-11.1 in the growth cone (Fig. 2h). Time-lapse recordings revealed that RAB-11.1 displayed a nearly identical movement pattern to that of GIP-2, with both proteins moving together in the growth cone (Fig. 2i, j). A frame-to-frame analysis of RAB-11.1 and GIP-2 distribution showed high correlation in the growth cone (Extended Data Fig. 2h), but no correlation in the cell body (Extended Data Fig. 2i)^19,20^. Interestingly, Rab11 itself and its associated endosomes have been found to influence the MT organization of mitotic spindles ^21^. We therefore interfered with RAB-11 function by expressing a dominant negative RAB-11.1 construct^22^, which caused a complete loss of GIP-2 puncta from the dendritic growth cone in about 35% of the animals (Fig. 2k, Extended Data Fig. 2j). Taken together, these results suggest that γ-TuRC localizes to RAB-11.1-positive endosomes in the growth cone to form the dgMTOC, which generates short PEO MTs in the distal growth cone and dynamic MEO MTs that populate the growing anterior dendrite.

This model predicts that the subcellular localization of the dgMTOC to the dendritic growth cone is instructive to populate the proximal dendrite with MEO MTs. We tested this model by perturbing the localization of the dgMTOC. We previously showed that the mature PVD anterior dendrite loses MEO MTs and gains PEO MTs in *unc-116/*kinesin-1 mutants^23^. Consistent with the MT polarity defect in mature PVD dendrites, *unc-116*(*e2310*) mutants showed a large decrease in the percentage of MEO MTs in the anterior dendrite shaft during outgrowth (93.8±2.7% (SEM, n=11)) in wt and 21.4±10.4% (SEM, n=12)) in *unc-116* mutants, Fig. 3a, Extended Data Fig. 3a). This dendritic MT polarity defect in *unc-116* mutants was also evident for GFP::TBA-1 dynamics (Extended Data Fig. 3b). Additionally, the reduced MEO frequency in *unc-116* mutants was partially rescued by expressing a wt *unc-116(+)* transgene during early neurite outgrowth (using the *unc-86* promoter), but not rescued by expressing *unc-116(+)* after anterior dendrite outgrowth (using a *ser-2* promoter), suggesting that UNC-116/Kinesin-1 is required specifically during early neurite outgrowth to establish dendritic MEO MTs (Extended Data Fig. 3a). Given that UNC-116 functions during an early developmental stage and that *unc-116* mutants display MT polarity defects during dendrite outgrowth, we considered that UNC-116 might be required for dgMTOC localization or function. To test this hypothesis, we examined EBP-2 dynamics in the outgrowing dendritic growth cone where the dgMTOC normally resides. In striking contrast to the wt controls, *unc-116* mutants lacked a convergence of PEO and MEO MTs at the outgrowing dendritic growth cone. Instead, the vast majority of MTs originated from the cell body and were PEO throughout the anterior dendrite (Fig. 3a, Extended Data Fig. 3c, d, Movie 4), raising the possibility that the dgMTOC is inactive or mislocalized in *unc-116* mutants. By tracing the origin of EBP-2 comets over time in *unc-116* mutants, we frequently observed that many EBP-2 comets emanated from a single region within the cell body (Extended Data Fig. 3e, Movie 5) – a stark contrast to the lack of a point-of-origin for EBP-2 comets in GIP-1^PVD(-)^ animals (Extended Data Movie 3). These results suggested that the dgMTOC remains in *unc-116* mutants but is mislocalized to the cell body, a possibility we assessed by examining the localization of GIP-2. Unlike in wt controls, GIP-2 was largely absent from the outgrowing dendrite growth cone and instead formed a cluster within the cell body in 64% (n=45) of *unc-116* mutants (Fig. 3b, Extended Data Fig. 3f). Occasionally, we found animals with a GIP-2 cluster in the posterior dendrite (Extended Data Fig. 3g), consistent with a rarely observed posterior dendrite-localized MTOC in *unc-116* mutants (Extended Data Fig. 3h). Together, these results indicate that the kinesin-1 motor is required for localizing the dgMTOC to the outgrowing dendritic growth cone. Furthermore, while depletion of GIP-1 in PVD showed that the dgMTOC is required to establish dendritic MEO MTs, this *unc-116* mutant phenotype suggests that the location of the dgMTOC instructs MT polarity.

**Fig. 3.**
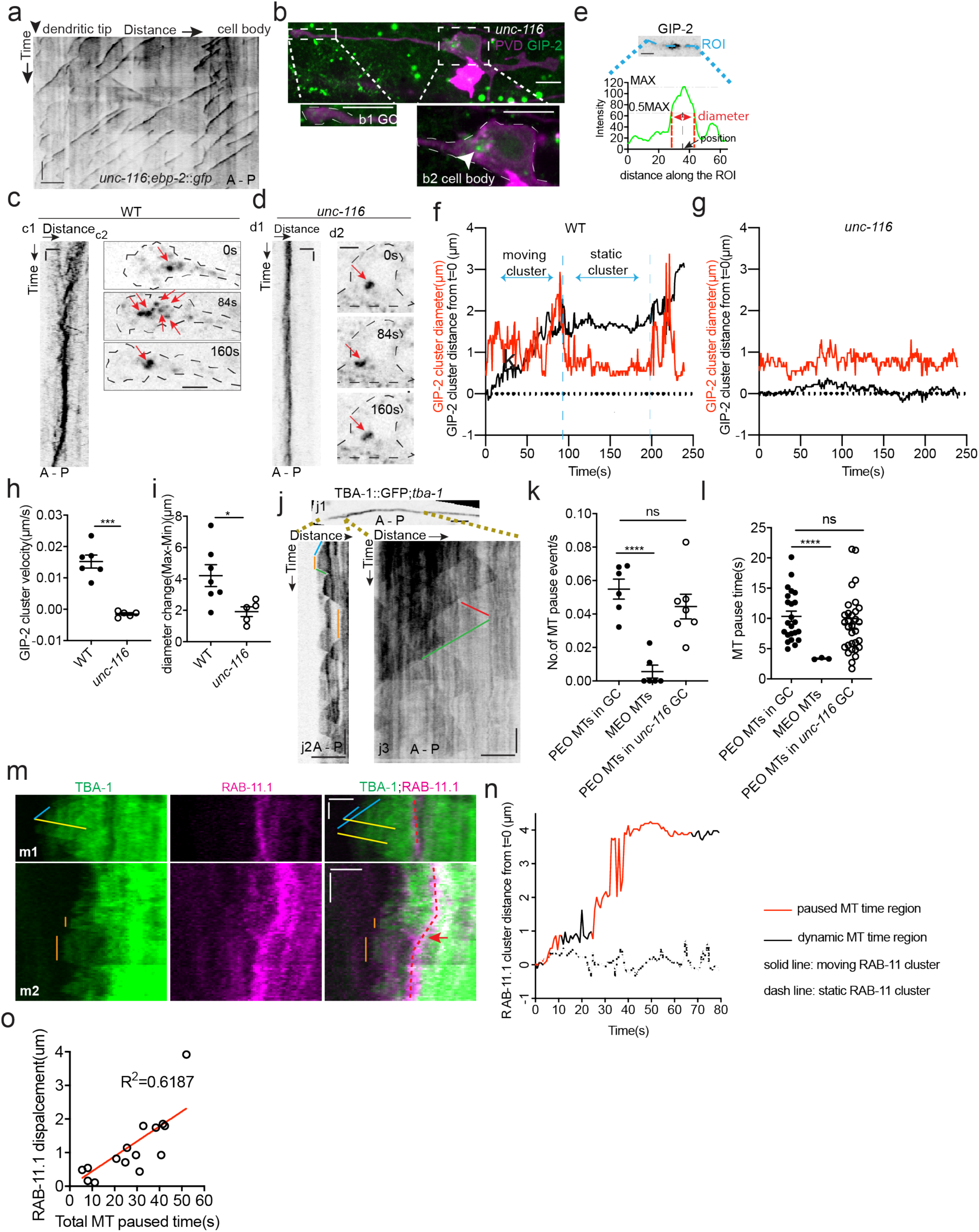
UNC-116/Kinesin-1 localizes the dgMTOC by moving it on transiently stable PEO MTs. (**a**) Kymograph of EBP-2::GFP in the outgrowing dendrite of an *unc-116* mutant. (**b**) GIP-2::GFP localization in PVD in an *unc-116* mutant. Growth cone (GC) region (b1); cell body region (b2); Arrowhead: GIP-2::GFP cluster in cell body. (**c-d**) GIP-2::GFP dynamics in the growth cone region. Wild type (c); *unc-116* mutant (d); kymograph of GIP-2::GFP (c1 and d1); GIP-2::GFP cluster in different time points (c2 and d2); dashed lines; outline of PVD growth cone (c2) and cell body (d2); red arrows; GIP-2::GFP clusters. (**e**) Schematic of GIP-2 diameter and position for quantification. Green line: GIP-2::GFP intensity along the blue ROI; red dashed lines: the edges of the GIP-2::GFP cluster used for diameter quantification; black arrow points to the black dashed line centered between the two red dashed lines, the position of GIP-2::GFP cluster. (**f-g**) GIP-2::GFP cluster diameter (red line) and distance from t=0 (black line) during GIP-2 cluster movement. Wild type (f); *unc-116* mutant (g). (**h)** Quantification of GIP-2::GFP cluster velocity in wt (n=6) and *unc-116* mutants (n=5). (**i**) Quantification of GIP-2 diameter change in wt (n=7) and *unc-116* mutants (n=5). (**j**) GFP::TBA-1 in the outgrowing dendrite (j1) and kymograph of GFP::TBA-1 in the indicated distal (j2) and proximal (j3) region. Blue line: growing PEO MTs in distal region; red line: growing MEO MTs in proximal region; green lines: retracting MTs; orange lines: pausing MTs. (**k-l**) Quantification of MT pause frequency (k) and pause time (i) of distal PEO and proximal MEO MTs. (**m**) Kymograph of GFP::TBA-1and mCherry::RAB-11.1 in the growth cone region. Blue line: growing PEO MTs; yellow lines: retracting MTs; orange lines: pausing MTs. Red dashed line indicates the position of mCherry::RAB-11.1 in the growth cone, red arrow indicates the moving mCherry::RAB-11.1 cluster. (**n**) Distribution of mCherry::RAB-11.1 cluster distance from t=0 correlated with MT dynamics over time. (**o**) Correlation of mCherry::RAB-11.1 displacement and distal PEO MT pause time (n=15). ***P<0.001, *P< 0.05, unpaired Student’s *t*-test, error bars represent SEM; A, anterior; P, posterior; vertical scale bar, 10s; horizontal scale bar, 2 μm in c, d, j and m, and 5 μm in others.

Next, we explored the mechanisms by which kinesin-1 localizes the dgMTOC. Time-lapse analyses of GIP-2 cluster movement showed that GIP-2 fluorescence undergoes processive movements towards the distal growth cone with several significant features. First, GIP-2 movements are largely unidirectional towards the distal dendrite tip, which is evident in kymograph analysis (c1 in Fig. 3c). Second, GIP-2 localization alternates between a tight cluster and a broader cluster with slightly more dispersed puncta during movement (c2 in Fig. 3c). Third, the movements are interspersed by pauses, which is evident by the alternating slopes and plateaus in GIP-2 displacement analyses (black trace in Fig. 3f, Extended Data Fig. 4a). Most GIP-2 exists in a bright cluster during pauses, while the fluorescence is dispersed into multiple smaller puncta during the mobile phase (Extended Data Movie 6). This phenomenon can be measured by the periodical changes of the diameter of the GIP-2 cluster during the movement cycle (Fig. 3e, red trace in Fig. 3f, Extended Data Fig. 4a). These results indicate that GIP-2 moves in a saltatory manner, with cycles of aggregation and dispersal, towards the distal dendritic tip. We hypothesized that kinesin-1 moves the GIP-2-positive endosomes along the PEO MTs in the growth cone. Consistent with this idea, *unc-116*/kinesin-1 mutants showed no displacement or dispersal of GIP-2 clusters (Fig. 3d, g-i, Extended Data Fig. 4b, Movie 7). These analyses suggest that UNC-116/kinesin-1 moves GIP-2 in the outgrowing dendrite.

With MTs emanating from the dgMTOC in both directions, the unidirectional movements of GIP-2 suggest that kinesin-1 prefers transport on the PEO MTs stretching towards the tip of the dendritic growth cone rather than the MEO MTs growing towards the cell body. To investigate differences between the populations of MTs originating from the dgMTOC, we examined TBA-1 dynamics in the outgrowing dendrite. Interestingly, we noticed that unlike MEO MTs which transitioned rapidly between polymerization and depolymerization (Fig. 2d, 2f, j3 in Fig. 3j, triangle shaped peaks in kymographs), the PEO MTs displayed frequent pauses between polymerization and depolymerization that lasted 10.34±0.85s (SEM, n=34) (j2 in Fig. 3j, plateaus, Fig. 3k, l), indicating that the distal PEO MTs are transiently stabilized, likely through interaction with the growth cone. We note that the distal PEO MTs originating closer to the cell body in *unc-116* mutants showed similar pause frequency and time as wt animals (Fig. 3k, l), indicating that the pausing of PEO MTs in the growth cone is independent of UNC-116 function. These data reveal an asymmetry in the behavior of dgMTOC-generated MTs, with PEO MTs pausing more than MEO MTs.

Kinesin-1 has been shown to prefer transporting cargoes on stable over dynamic MTs^24–26^. To assess whether a preference of kinesin-1 for stable MTs could explain the biased dgMTOC movements, we simultaneously recorded TBA-1 dynamics and RAB-11.1 movements and assessed whether endosome movement corresponded with the presence of paused MTs and not with dynamic MTs. No obvious distal directed RAB-11.1 movements were observed when only dynamic PEO MTs are present (m1 in Fig. 3m). In contrast, endosome movements toward the distal dendritic tip were observed in the presence of stable PEO MTs (m2 in Fig. 3m, orange lines). To correlate the RAB-11.1 movements with MT dynamics, we quantified the displacement of RAB-11.1 over time and determined and labeled time periods in which a PEO MT was paused (red, Fig. 3n) and periods in which PEO MTs were only dynamic (black, Fig 3n). We found that endosome movements preferentially occurred during periods with paused MTs (solid line in Fig. 3n) and there was no obvious RAB-11.1 movement when the PEO MTs were dynamic (dashed line in Fig. 3n). By sampling of a population of worms, we found that RAB-11.1 endosome displacement showed a positive correlation with the total MT pause time overall (Fig. 3o). These results suggest that distally-directed endosomes move on paused or stable PEO MTs in a kinesin-1-dependant manner.

Taken together, these findings are consistent with a model in which kinesin-1 transports γ-TuRC positive endosomes towards the dendritic tip as the growth cone advances. The endosomes act as the dgMTOC to build PEO MTs towards the growth cone, which in turn function as MT tracks for the next bouts of kinesin-1-mediated endosome movements. The PEO MTs are stable for about 10s, which is long enough to support the MTOC movements, but also are transient in nature as these MTs showed depolymerization events after pausing (j2 in Fig. 3j). We speculate that the transient nature of these “stabilized” PEO MTs likely ensures that the majority of MTs are still MEO once the MTOC has passed this area.

While kinesin-1-mediated motility explains the processive movements of the dgMTOC towards the dendritic tip, it is not clear how the multiple γ-TuRC puncta aggregate to re-form a single cluster after its dispersal during movement, which is likely critical to maintain a singular dgMTOC. Cytoplasmic dynein has been shown to concentrate organelles such as Golgi or RAB-11 endosomes near MTOCs in non-neuronal cells^27,28^. We therefore tested whether dynein is required for dgMTOC clustering by examining *dhc-1*/dynein heavy chain mutants. *dhc-1(or195)*, a temperature sensitive dynein mutant, displayed several types of defects in dgMTOC localization when animals were cultured at nonpermissive temperature. In ∼30% of *dhc-1* mutants, GIP-2 clusters were completely absent from the PVD anterior dendrite and the rest of the cell (compare wt localization in Fig. 4a to b1 in Fig. 4b, Fig. 4c). In another ∼40% of mutant animals, one or several GIP-2 cluster(s) could be found in the cell body and occasionally in the posterior dendrite (b2 in Fig. 4b, Fig. 4c). In the remaining ∼30% of mutant animals, a GIP-2 cluster still localized to the anterior dendrite but showed a variable distance to the dendritic tip which was longer than in wt animals (b3 in Fig. 4b, Fig. 4c, d). Time-lapse imaging analysis of GIP-2 movements in the subset of *dhc-1* mutant worms which have an anterior dendrite-localized GIP-2 cluster showed that net GIP-2 movement towards the dendritic tip was reduced compared to that of wt, consistent with its ectopic localization (Fig. 4e-i, Extended Data Fig. 4c). Strikingly, further analyses of *dhc-1* mutants showed that GIP-2 puncta exhibited excessive, continuous movements with constant dispersal but failed to form a relatively stable singular cluster (Fig. 4f, h). The continuous dispersal of GIP-2 clusters lead to a reduction in the maximal intensity of the cluster over time which might explain the lack of GIP-2 clusters in a subset of *dhc-1* mutants (Fig. 4j, k). Endogenously-tagged DHC-1 localized to the dgMTOC region as a cluster and like the GIP-2 clusters remained associated with the base of the advancing growth cone (Fig. 4l). These dynamic movements closely coincided with those of RAB-11.1 foci (Fig. 4m). This striking subcellular localization of DHC-1 further suggests that DHC-1 functions locally at the dgMTOC. Consistent with the defect in dgMTOC localization, we found that about 50% of *dhc-1* mutants showed complete reversal of MT polarity while another 20% of mutants showed mixed PEO and MEO MTs in the anterior dendrites perhaps due to the dispersed GIP-2 clusters in these mutants (Fig. 4n, o). These results suggest that dynein uses its minus-end-directed motor activity to re-cluster γ-TuRC-positive endosomes together which had dispersed during translocation towards the distal tip of the dendritic growth cone by kinesin-1.

**Fig. 4.**
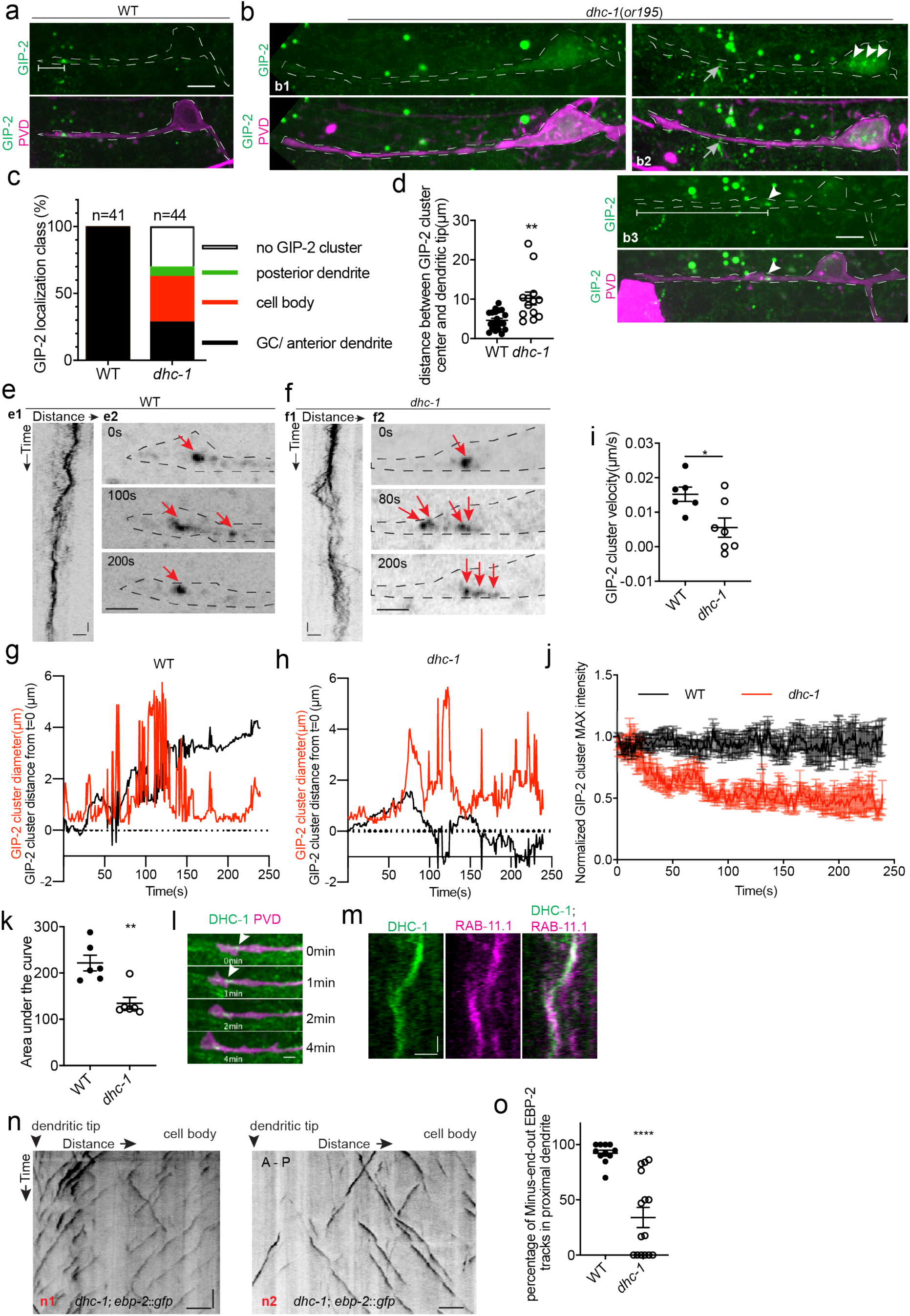
DHC-1/dynein clusters GIP-2 to form a single dgMTOC during outgrowth. (**a-b**) Endogenous GIP-2::GFP localization in wild type (a) and *dhc-1* mutants (b). White arrowheads, GIP-2::GFP clusters; gray arrows in b2, unrelated signal from gut granules; white brackets, distance between GIP-2 cluster and dendritic tip. (**c**) Quantification of GIP-2::GFP class of localization in *dhc-1* mutants. (**d**) Quantification of distance between GIP-2::GFP cluster and dendritic tip in wt (n=20) and *dhc-1* mutants (n=13). (**e-f**) GIP-2::GFP dynamics in wt (e) and *dhc-1* mutants (f). Kymograph of GIP-2::GFP (e1 and f1); GIP-2::GFP cluster at different time points (e2 and f2); dashed lines, outlines of PVD dendrite; red arrows, GIP-2::GFP clusters. (**g-h**) Distribution of GIP-2::GFP cluster diameter (red line) and distance from t=0 (black line) during GIP-2::GFP cluster movement in wt (g) and *dhc-1* mutant (h). (**i**) Quantification of GIP-2::GFP cluster velocity in wt (n=6) and *dhc-1* mutants (n=7). (**j**) Quantification of GIP-2::GFP cluster maximum intensity in wt (n=7, black) and *dhc-1* mutants (n=7, red). (**k**) Quantification of the area under the lines in **j**. (**l**) Endogenous DHC-1::GFP localization in an outgrowing dendrite at different time points. White arrowheads, DHC-1::GFP in the growth cone. (**m**) Kymograph of DHC-1::GFP and mCherry::RAB-11.1 in the growth cone region. (**n**) Kymograph of EBP-2::GFP in a *dhc-1* outgrowing anterior dendrite. (**o**) Quantification of MT polarity in wt (n=11) or *dhc-1* mutants (n=15). Horizontal scale bar, 10s. Vertical scale bar, 2 μm for e1, f1 and m; others, 5 μm. ****P<0.0001, **P<0.01, *P< 0.05, unpaired t test, error bars represent SEM.

Collectively, these data support a model in which MTs in the dendrite are locally generated by a γ-TuRC-positive endosome-based dgMTOC which advances with the growing tip of the dendrite by the concerted action of both kinesin-1 and dynein (Extended Data Fig. 5). This dgMTOC generates PEO MTs that extend towards the growing dendrite tip and MEO MTs that grow towards the cell body. The PEO MTs are transiently stabilized by interaction with the growth cone and serve as the tracks on which UNC-116/kinesin-1 preferentially transports the γ-TuRC-bearing endosomes into the anterior dendrite as it emerges, and further towards the distal dendritic tip as it grows. These transport events disperse the endosomes, which are refocused by dynein-mediated minus-end-directed movements to maintain a single MTOC throughout dendrite outgrowth. Dynein-based clustering and kinesin-based movement are intermixed, leading to interspersed anterograde movement and dispersal and re-clustering of the dgMTOC.

Importantly, our data demonstrate that the MEO orientation of MTs in the dendrite – which is essential for their function in polarized trafficking – is established by a remarkably mobile MTOC generated on endosomes. Endosomes in mitotic cells can deliver spindle material to the centrosome, suggesting that endosomes might be a general source of MTOC material ^21^. Significantly, this mobile MTOC instructs the unique MEO MT dendritic polarity from the earliest stages of dendrite outgrowth by generating numerous MEO MTs which populate the dendrite as the growth cone advances. While only present during dendrite outgrowth, this MTOC is responsible for establishing the polarized MT orientation seen in mature dendrites. We speculate that a small percentage of the MEO MTs generated by the dgMTOC are ultimately stabilized to form the MEO MTs in mature dendrites. In mammalian neurons, dendrites contain both PEO and MEO MTs. We imagine that diverse cell types can tune their MT dynamics to achieve different relative levels of PEO and MEO MTs to generate and maintain a mixed polarity dendritic MT population in combination with the activity of a growth cone localized MTOC.

**Table 1.** *C. elegans* strains used in this study.

| Strain | Genotype |
| --- | --- |
| TV24185 | *zif-1(gk117)* III; *wyEx9745*[*Punc-86::mCherry::PLCdeltaPH*] |
| TV24455 | *zif-1(gk117)* *gip-1(wow5[zf::gfp::gip-1])* III; *wyEx9745* |
| TV24458 | *ebp-2(wow47[ebp-2::gfp::3xflag])* II; *zif-1(gk117)* III; *wyEx9745* |
| TV24424 | *gip-2(lt19[gip-2::gfp::loxP::cb-unc-119(+):loxP])* I; *wyEx9745* |
| TV24830 | *ebp-2(wow47)* II; *gip-1(wow25[tagRFP-t::3xMyc::gip-1])* III; *wyEx9745* |
| TV24645 | *zif-1(gk117)* *gip-1(wow5[zf::gfp::gip-1])* III; *wyEx9744*[*Punc-86::mCherry::PLCdeltaPH Punc-86::zif-1*] |
| TV24646 | *ebp-2(wow47[ebp-2::gfp::3xflag])* II; *zif-1(gk117)* *gip-1(wow5[zf::gfp::gip-1])* III; *wyEx9744*[*Punc-86::mCherry::PLCdeltaPH Punc-86::zif-1*] |
| TV25492 | *gip-2(lt19)* I; *wyIs581[ser-2P3::myri-mCherry]* |
| TV21720 | *tba-1(ok1135)* II; *wyIs813*[*Punc-86::gfp::tba-1 Punc-86::mCherry::PLCdeltaPH*] |
| TV25234 | *gip-2(lt19)* I; *glo-1(zu391)* X; *wyEx10042*[*Punc-86::mCherry::rab-11.1 cDNA Podr-1::gfp*] |
| TV25235 | *gip-2(lt19)* I; *glo-1(zu391)* X; *wyEx10042*[*Punc-86::mCherry::RAB-11.1(S25N) Podr-1::gfp*] |
| TV24622 | *wyEx9876* [*Punc-86::AMAN-2(1-84aa)::GFP novo2 Punc-86::mCherry::PLCdeltaPH Podr-1::gfp*] |
| TV25150 | *wyEx10015*[*Punc-86::gfp::rab-6.2 Punc-86::mCherry::rer-1 Podr-1::gfp*] |
| TV25463 | *glo-1(zu391)* X; *wyEx10092*[*Punc-86::gfp::rab-11.1 cDNA Punc-86::mCherry::rab-6.2 Podr-1::gfp*] |
| TV23841 | *ebp-2(wow47)* II; *unc-116(e2310)* III |
| TV24433 | *ebp-2(wow47)* II; *unc-116(e2310)* III; *wyEx9745* |
| TV24434 | *gip-2(lt19)* I; *unc-116(e2310)* III; *wyEx9745* |
| TV21585 | *tba-1(ok1135)* I; *unc-116(e2310)* III; *wyEx8784*[*Punc-86::gfp::tba-1 Punc-86::mCherry::PLCdeltaPH*] |
| TV25470 | *tba-1(ok1135)* I; *wyIs813*; *wyEx10099*[*Punc-86::mcherry::rab-11.1 cDNA Podr-1::gfp*] |
| TV25536 | *dhc-1(or195)* *gip-2(lt19)* I; *wyEx9745* |
| TV24721 | *dhc-1(ie28 [dhc-1::degron::gfp])* I; *wyEx9745* |
| TV25507 | *dhc-1(ie28)* I; *glo-1(zu391)* X; *wyEx10110* [*Punc-86::mCherry::RAB-11.1 cDNA Podr-1::gfp*] |
| TV25111 | *dhc-1(or195)* I; *ebp-2(wow47)* II; *wyEx9745* |
| TV25113 | *gip-2(lt19)* I; *lin-5(ev571)* II; *wyEx9745* |
| TV25054 | *lin-5(cp288)* II; *wyEx9745* |
| TV201 | *wyIs22*[*Punc-86::rab-3a::gfp Podr-1::rfp*] |

**Table 2.**
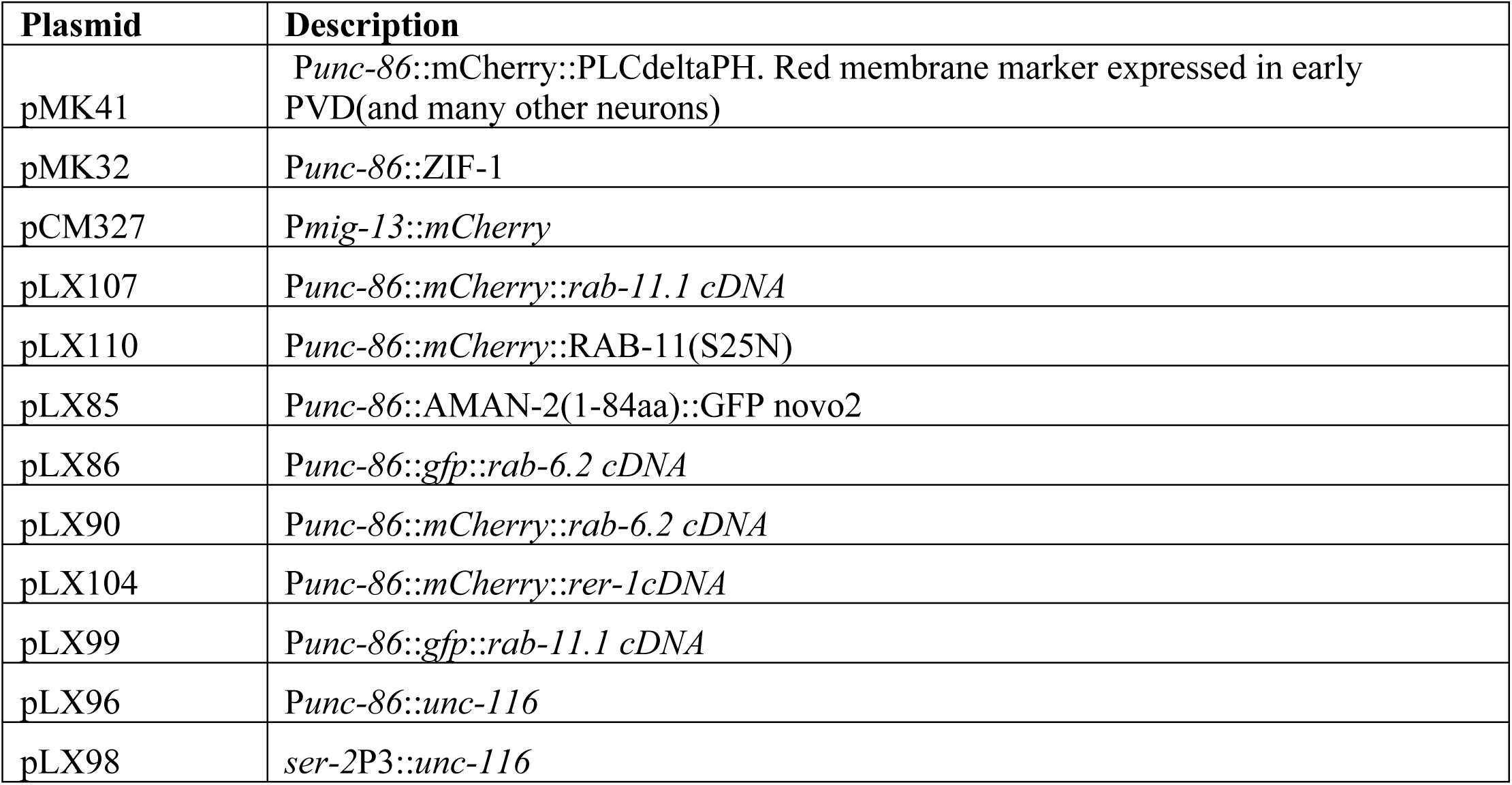
Plasmids used in this study.

**Table 3.**
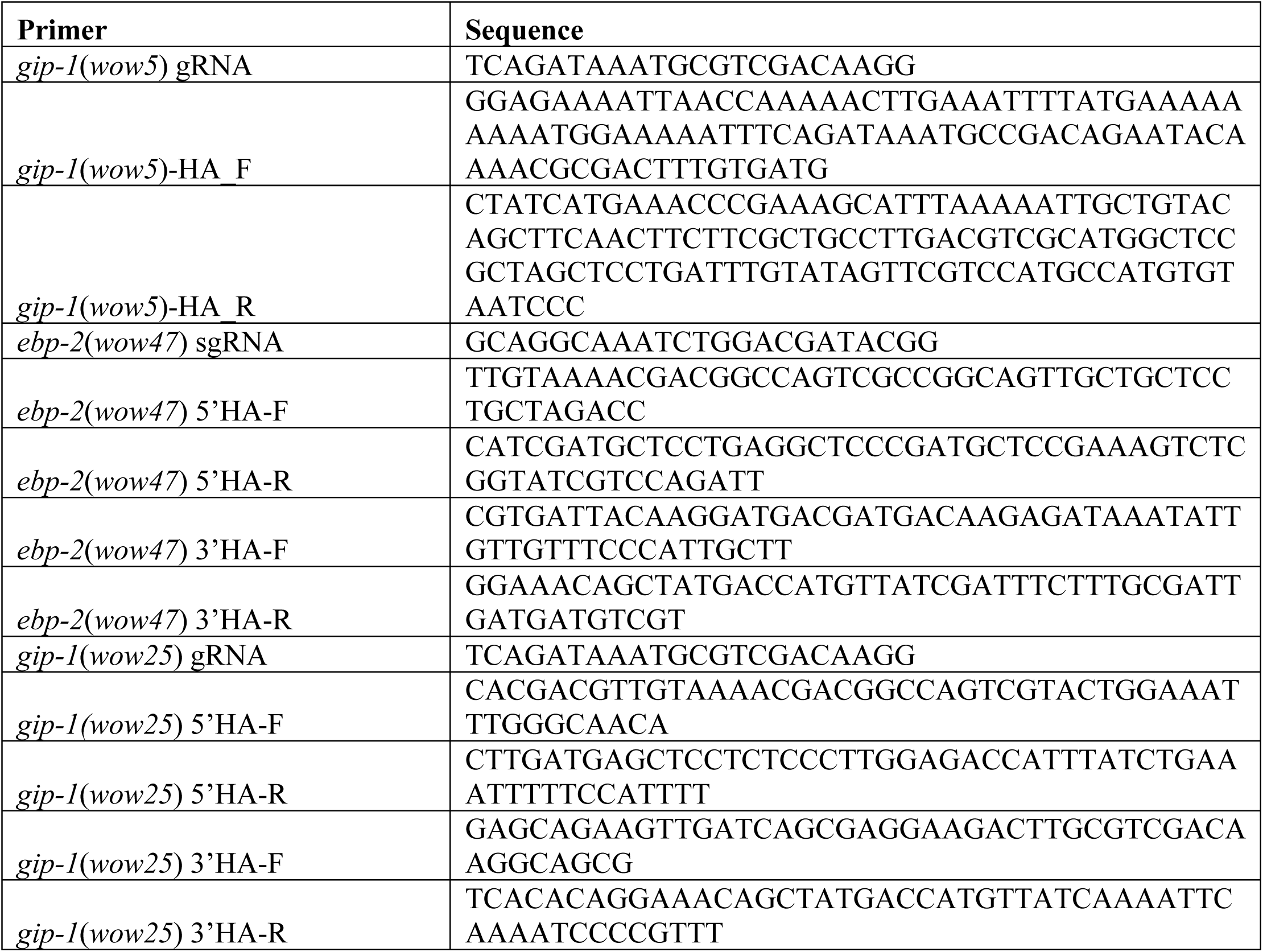
Primers used in this study.

## Acknowledgments

We thank members of the Shen lab for their scientific feedback and discussion.

## Author contributions

K. Shen, X. Liang, M. Kokes, J. Feldman and A. Moore conceptualized aspects of the study. X. Liang, M. Kokes, R. Fetter and A. Moore performed key experiments and analyzed data. X. Liang, M. Kokes, M. Pickett, M. Sallee, and A. Moore made reagents and strains. K. Shen, X. Liang, M. Kokes, J. Feldman and A. Moore wrote the manuscript and made the figures.

## Data and materials availability

All primary data is available upon request.

---

**Extended Data Fig. 1.**
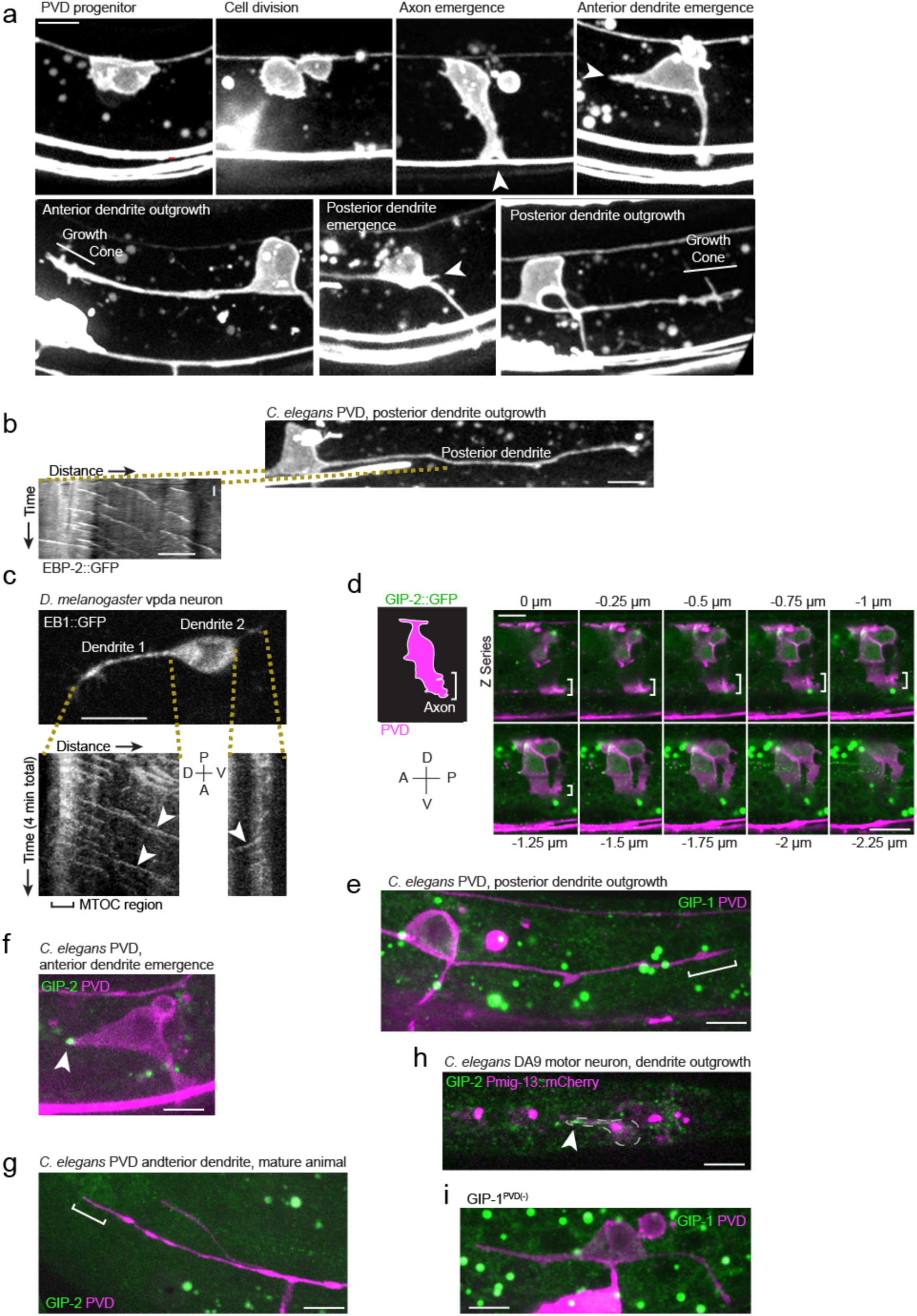
The dgMTOC is unique to outgrowing anterior dendrites and is conserved in *C. elegans* and *D. melanogaster*. (**a**) Early PVD morphogenesis events occur with a stereotyped temporal sequence and orientation, arrowheads: site of emerging neurite (axon, anterior dendrite, posterior dendrite). Unrelated fluorescence: horizontal lines at bottom of images (other neurons), bright spots (gut granules). (**b**) Morphology of PVD during posterior dendrite outgrowth (top right) and kymograph (bottom left) of EBP-2::GFP in indicated region, vertical scale bar 10s. Note that the posterior dendrite behaves differently from the anterior dendrite - it outgrows later and has majority PEO MTs (**c**) Top: A *Drosophila* vpda neuron expressing EB1::GFP in the early stage of dendrite outgrowth with two dendritic processes emanating from the cell body. Bottom: Kymographs of EB1::GFP in each process over four minutes. White arrowheads: EB1 comets generated from a region near the tip of the dendrite and moving towards the cell body. (**d**) Lateral view of PVD through three-dimensional space. Cartoon at left drawn from a maximum projection of the cell depicts the boundaries of PVD. The outgrowing axon is indicated with white brackets. Because structures outside the cell can appear to be inside following z-slice projection, serial z-stack images from the top to the bottom of the cell are displayed as a montage to the right. Note the absence of endogenously-tagged GIP-2::GFP (green) in the outgrowing PVD axon (white brackets). (**e**) GFP::GIP-1 (green) localization in the outgrowing posterior PVD dendrite (magenta). White brackets: growth cone. Note the absence of a GIP-1 cluster. (**f**) GIP-2::GFP localization in the emerging anterior PVD dendrite. White arrowhead: GIP-2::GFP cluster. (**g**) GIP-2::GFP localization in mature dendrite. White bracket: distal dendrite. (**h**) GIP-2::GFP localization in an outgrowing *C. elegans* DA9 dendrite. (**i**) GFP::GIP-1 localization in a GIP-1^PVD(-)^ animal. All images are oriented with anterior to the left and posterior to the right; unrelated structures within *C. elegans* tissues include gut granules (large green dots); horizontal scale bars, 5μm.

**Extended Data Fig. 2.**
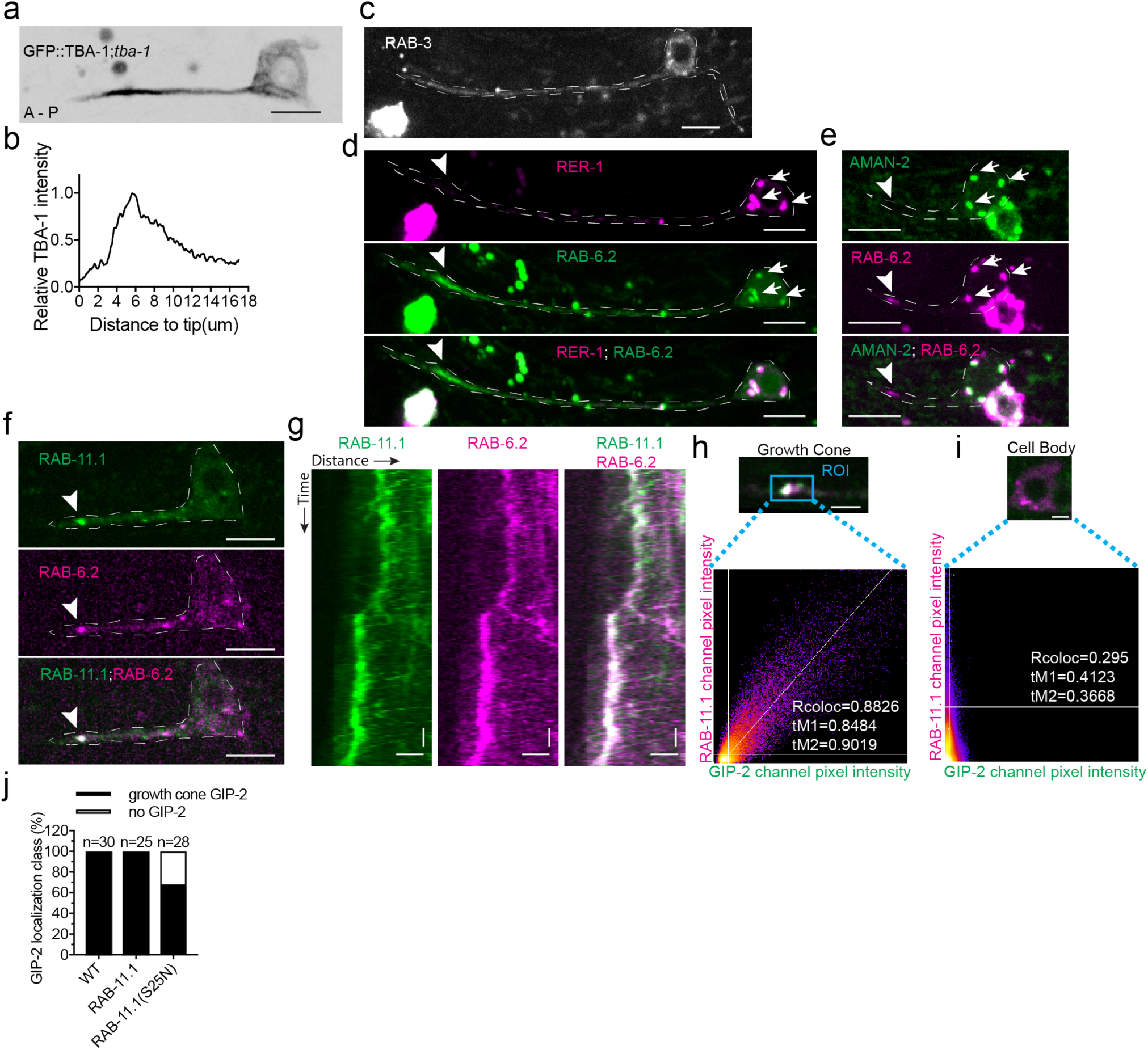
Golgi and synaptic vesicles do not localize to the growth cone region. (**a-b**) GFP::TBA-1 in an outgrowing dendrite (**a**), Distribution of relative TBA-1 intensity from the dendritic tip to the cell body(**b**). (**c-e**) GFP::RAB-3 (c), mCherry::RER-1 and GFP::RAB-6.2 (d), and AMAN-2::GFP and mCherry::RAB-6.2 (e) localization in outgrowing anterior PVD dendrite. White arrows: Golgi stack in cell body; white arrowheads: RAB-6.2 in growth cone. (**f**) GFP::RAB-11.1 and mCherry::RAB-6.2 localization in outgrowing dendrite. White arrowhead: GFP::RAB-11.1 and mCherry::RAB-6.2 in growth cone. (**g**) Kymograph of GFP::RAB-11.1 and mCherry::RAB-6.2 in growth cone region. Horizontal scale bar, 2 μm; vertical scale bar, 10s. (**h-i**) Frame to frame colocalization analysis between GIP-2::GFP and mCherry::RAB-11.1 in growth cone (h) and cell body (i). (**j**) Quantification of GIP-2::GFP class of localization in wt worms or worms overexpressing a RAB-11 or RAB-11(S25N) dominant negative mutant. All images: Scale bar, 5 μm (except g); white dashed line, outline of PVD; anterior to the left and posterior to the right. Images in f were taken in the *glo-1*(*zu391*) mutant background to reduce the gut granule signal.

**Extended Data Fig. 3.**
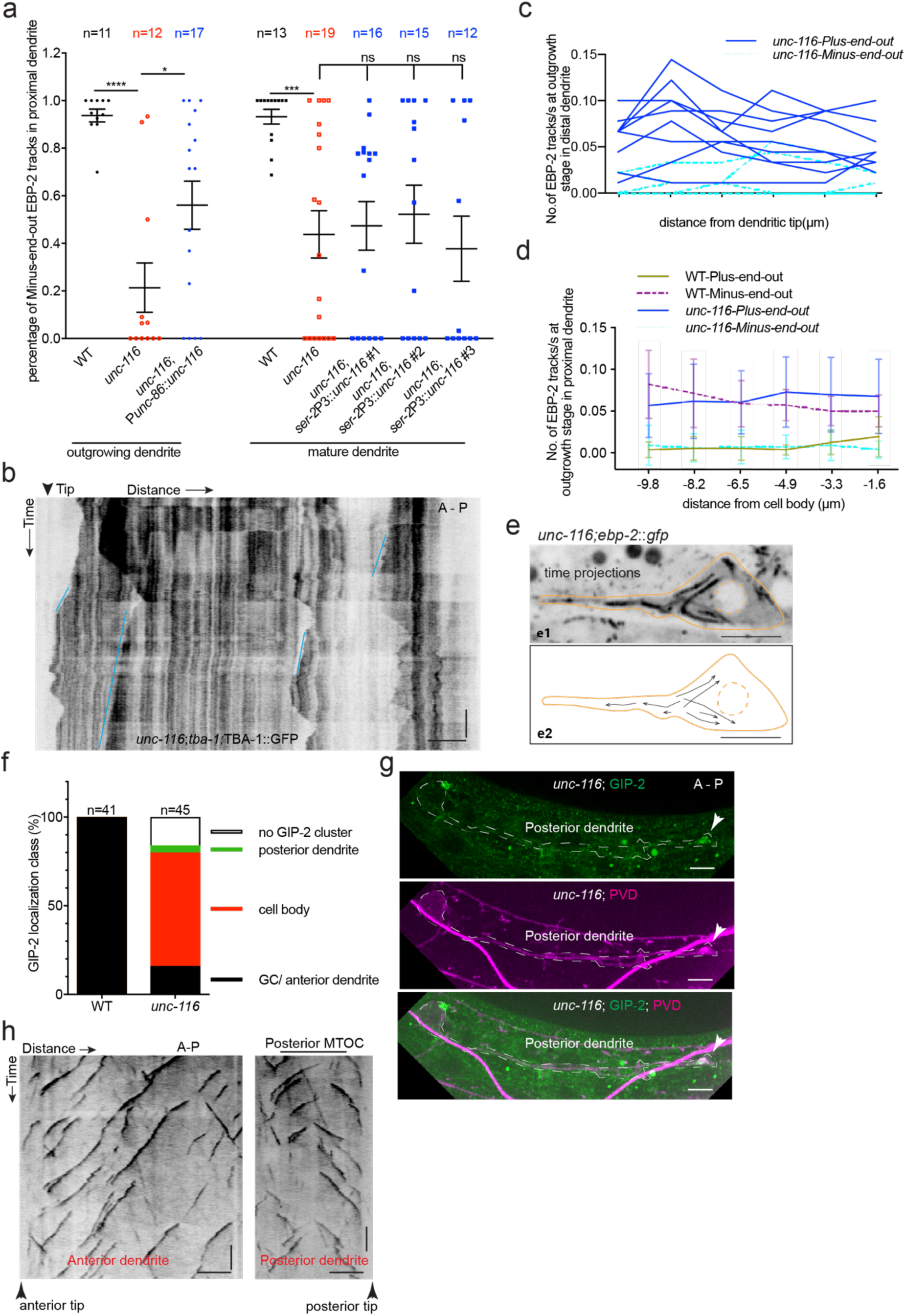
MT polarity and dgMTOC localization require UNC-116/kinesin-1. (**a**) Quantification of MT polarity in wt, *unc-116* mutants, and *unc-116* mutants expressing UNC-116 early (*unc-86* promoter), and late (*ser-2* promoter). (**b**) Kymograph of GFP::TBA-1 in an *unc-116* mutant. Blue lines: PEO MTs. (**c-d**) Plus-end-out (PEO) and minus-end-out (MEO) MT polymerization frequency measured by EBP-2::GFP tracks at interval distances from the dendritic tip towards the cell body (c) in *unc-116* mutants (n=9) and from the cell body outwards towards the dendritic tip (d) in both WT (n=7) and *unc-116* mutants (n=11). Distances on the x-axis are displayed to imitate the anterior-left and posterior-right orientation. (e)Top (e1): Time projection of EBP-2::GFP dynamics in an *unc-116* mutant. Bottom (e2): Schematic of the EBP-2::GFP trajectories generated by tracing the EBP-2::GFP frame by frame. Orange line: the outline of the outgrowing PVD; dashed orange circle: nucleus; black arrows: EBP-2::GFP trajectories.(**f**) Quantification of GIP-2::GFP class of localization in wt and *unc-116* mutants. (**g**) Posterior dendrite localized GIP-2::GFP in an *unc-116* mutant. White arrowhead, GIP-2::GFP in growth cone. (**h**) Kymograph of EBP-2::GFP in both anterior (left) and posterior (right) dendrite in *unc-116* mutant which showed a posterior dendrite localized MTOC. Horizontal scale bar, 5 μm; vertical scale bar, 10s; ****P<0.0001, ***P<0.001, *P<0.5, unpaired Student’s *t*-test, error bars represent SEM.

**Extended Data Fig. 4.**
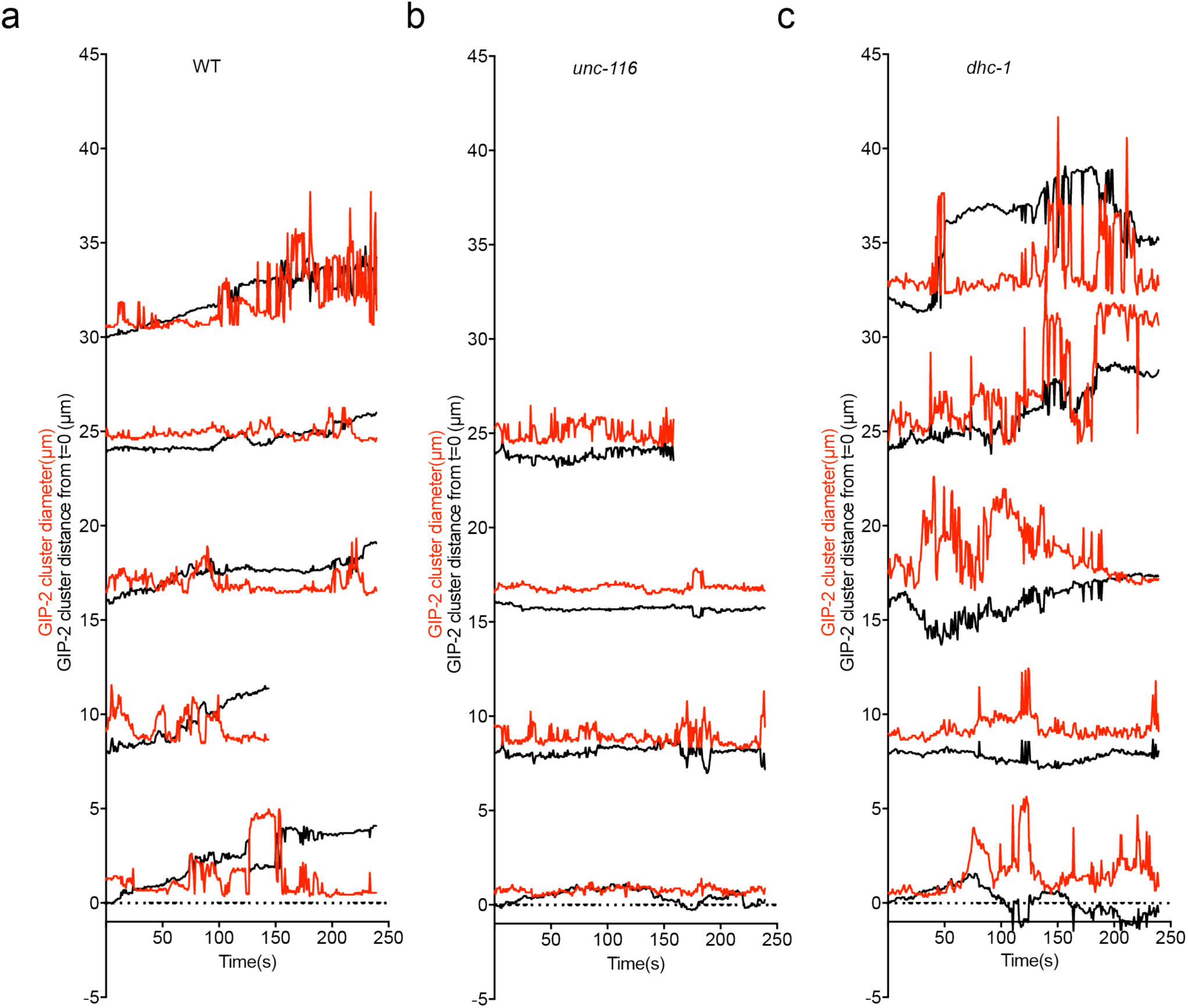
Distribution of GIP-2::GFP cluster distance from t=0 (black line) and diameter (red line) during GIP-2 cluster movement in different wt (n=5, a), *unc-116* mutants (n=4, b), and *dhc-1* mutants (n=5, c).

**Extended Data Fig. 5.**
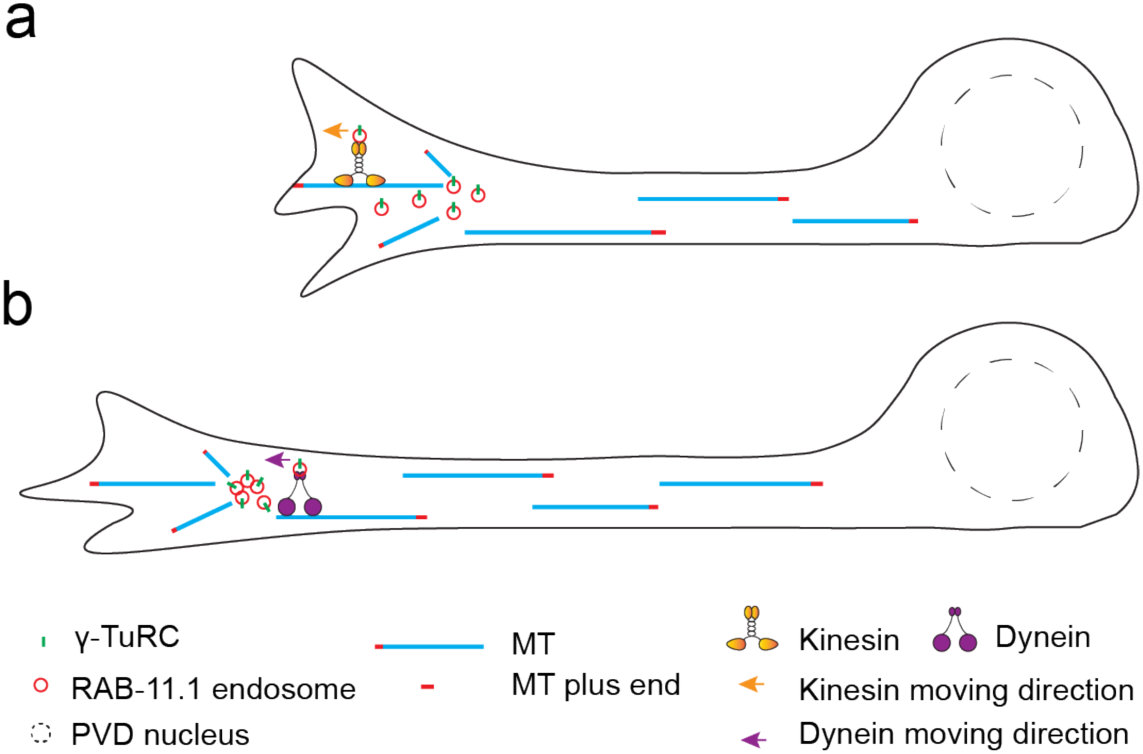
Proposed model for the movement of the endosome-based dgMTOC in the growth cone region. (**a**) The dgMTOC generates PEO MTs that extend towards the dendrite tip and MEO MTs that grow towards the cell body. The PEO MTs are transiently stabilized by interaction with the growth cone and serve as the tracks on which kinesin-1 transports the endosomes further towards the distal dendritic tip. (**b**) Dynein clusters the endosomes to form a single MTOC and maintain the structure stay in the growth cone in between the bouts of kinesin mediated movements.

## Materials and Methods

### *C. elegans* strains

Worms were raised on NGM plates at 20°C using OP50 *Escherichia coli* as a food source. Worm strains which contain temperature sensitive mutant *dhc-1*(*or195*) were maintained in 16°C and were shift to 25°C for phenotype analysis. *C. elegans* strains used in this study are listed in table S1. OD2509 [*gip-2*(*lt19*[*gip-2::gfp*]::*loxP::cb-unc-119(+*)::*loxP*) I; *unc-119(ed3)* III] was a gift from Dr. Karen Oegema at the University of California San Diego^36^. The generation of JLF273 *ebp-2(wow47[ebp-2::gfp]) II; zif-1(gk117) III; wowEx10* (used to make TV23841), JLF38 *gip-1(wow5[zf::gfp::gip-1]) zif-1(gk117) III; wowEx10* (used to make TV24455), and JLF155 *zif-1(gk117)* has been described^37^.

### Electron Microscopy

Worms were prepared for conventional EM by high pressure freezing/freeze-substitution. Worms in E. coli containing 20% BSA were frozen in 100 µm well specimen carriers (Type A) opposite a hexadecane coated flat carrier (Type B) using a BalTec HPM 01 high pressure freezer (BalTec, Lichtenstein). Freeze-substitution in 1% OsO4, 0.1% uranyl acetate, 1% methanol in acetone, containing 3% water^29,30^ was carried out with a Leica AFS2 unit. Following substitution, samples were rinsed in acetone, infiltrated and then polymerized in Eponate 12 resin (Ted Pella, Inc., Redding, CA). Serial 50 nm sections were cut with a Leica UCT ultramicrotome using a Diatome diamond knife, picked up on Pioloform coated slot grids and stained with uranyl acetate and Sato’s lead^31^. Sections were imaged with an FEI Tecnai T12 TEM at 120 kV using a Gatan 4k x 4k camera. TrakEM2 in Fiji was used to align serial sections^32,33^. Modeling of serial sections was performed with IMOD^34^.

### Molecular biology, transgenic lines, and CRISPR

Plasmids and primers used to generate transgenic or knock in *C. elegans* strains in this study are listed in tables S2 and S3. Expression clones were made in the pSM vector, a derivative of pPD49.26 (A. Fire) with extra cloning sites (S. McCarroll and C.I. Bargmann, personal communication). Plasmids and sequences are available upon request. P*unc-86*::mCherry::PLCdeltaPH was generated using Gibson cloning. Transgenic strains (1–50 ng/μl) were generated using standard techniques and coinjected with markers P*odr-1*::GFP or P*odr-1::*RFP.

### *C. elegans* synchronization and staging

Gravid adults were bleached in a hypochlorite solution to obtain embryos which were washed in M9 and allowed to hatch either in M9 or on unseeded NGM plates overnight to obtain a population synchronized in L1 arrest. This L1-arrested population was kept for a maximum of 5 days and transferred to OP50-seeded NGM plates at different times to achieve specifically aged synchronized L2 populations for imaging.

To image early outgrowing PVD neurites, wild type and GIP-1 knockdown L1-arrested animals were grown on OP50-seeded NGM plates at 22°C for 18-19 hours or 25°C for 16-17 hours before imaging. *unc-116*(*e2310*) L1-arrested animals were grown on OP50-seeded NGM plates at 22°C for 20-22 hours or 25°C for ∼18 hours before imaging.

*dhc-1* L1-arrested worms were obtained at 25°C and then were grown on OP50 seeded NGM plates at 25°C for 16-18 hours.

Wild type GFP::TBA-1 animals were imaged 1.5-2 hours later than other wild type image to get a more clear MT dynamic kymograph in the growth cone region, as in the later outgrowth stage, the MT number in the growth cone region will slightly reduced.

To image mature PVD neurites, bleached adults were placed on OP50-seeded NGM plates for about 48 hours at 20°C, then the mid L4 worms were picked to image for EBP-2 dynamics and GIP-2 localization.

To image the outgrowing DA9 neuron, some 3-fold stage embryos were transferred to a new plate and then the L1 worms right after hatching were imaged.

### Slide preparation

Just prior to imaging, *C. elegans* animals were mounted on 3-5% agarose pads. L2s or L1s were picked and released into a 1 μl M9 or water droplet on an inverted coverslip while minimizing bacterial transfer. Prior to mounting on the freshly made agarose pad, the droplet was surrounded by a 1 μl droplet of 0.05 µm Polysterene Polybeads (Polysciences) and a droplet of levamisole (final concentration of approximately 3 mM) to immobilize worms. L4s were picked to M9 directly on the agarose pad with levamisole added prior to mounting. Slides were sealed with VALAP or Vaseline prior to time-lapse imaging. All imaging was performed within 40 minutes of mounting.

### Microscope system

Imaging of *C. elegans* was performed on an inverted Zeiss Axio Observer Z1 microscope equipped with a Yokogawa spinning disk, QuantEM:512SC Hamamatsu camera (set to 600 EM Gain), a Plan-Apochromat 100x/1.4 NA objective (Zeiss), 488 nm and 561 nm lasers, and controlled by MetaMorph Microscopy software (Molecular Devices).

### Imaging parameters

Endogenous EBP-2::GFP dynamics in outgrowing and mature PVD dendrites were imaged using 50% 488 nm laser power, 100 ms exposure, and with 200 ms time interval between acquisitions. 200-300 frames were recorded per animal. The same imaging parameters were also used to image GFP::TBA-1 dynamics in outgrowing dendrite.

To assess protein localization within PVD during neurite outgrowth, endogenously-GFP-tagged GIP-1, GIP-2 were imaged using 70-80% 488 nm laser power recorded with 300 ms exposure. The membrane mCherry co-marker was imaged using 70%-80% 561 nm laser power with 200 ms exposure. To fully sample PVD in three dimensions, z-sections were imaged at Nyquist resolution, every 0.25 μm from above to below a single PVD in each animal. Micrographs in Fig. 1c, and Extended Data Fig. 1d are maximum projections of the z-sections that include the PVD dendrite.

For time-lapse imaging of these strains to assess dynamics of complex localization to the outgrowing dendrite tip, 488 nm laser power was reduced to 50% with 200 ms exposures to reduce phototoxicity and photobleaching. Up to 4 z-sections were imaged at 0.5 μm, and acquisitions occurred every 15 or 30 seconds for up to a total of 60 minutes. Note that Fig. 1d displays only frames every 60 seconds even if animals were imaged more frequently. Only animals which displayed a steady rate of growth cone advance indicating healthy animals were used for analysis.

For time lapse imaging to look at the details of GIP-2, RAB-11.1 and DHC-1 dynamics at the growth cone region, only one z section is acquired, and images were taken every 200ms for up to 200-300 frames, and images were taken with a 100ms exposure time and 70% laser power for both 488 nm and 561 nm channels. To image TBA-1 together with RAB-11.1, the 488 nm laser power was reduced to 50% while other parameters were the same.

### Image analysis and quantification

Images were processed and analyzed using MetaMorph (Molecular Devices) and ImageJ to create kymographs or psuedocolored merged maximum intensity micrographs and assembled into figures using Adobe Photoshop and Illustrator. Statistical calculations and graphing were done in Prism 7 (GraphPad).

To display the dynamics of endogenously-GFP-tagged GIP-1 and GIP-2 localization to the PVD dendritic growth cone over time in Fig. 1d, a 20 pixel-wide line segment was drawn over the growth cone outgrowth region and processed using the ImageJ function ‘Straighten’ to slightly straighten the region. The Make Montage function was then used to create a montage displaying that region every 60 seconds. Due to minor movements of the worm, uncropped merged time-lapse images were registered to each other using the ImageJ StackReg function before making a montage for GIP-1.

Kymographs were made in ImageJ by drawing a straight or segmented line (width of 6-10 pixels) along a process and using the KymographBuilder plugin or Stacks-Reslice function on the stacked time-lapse file.

To quantify EBP-2::GFP comet direction and frequency at interval distances across the outgrowing dendrite (line graphs in Fig. 1b, Extended Data Fig. 3c, d), kymographs were created from recordings of numerous animals and the number of EBP-2::GFP MEO and PEO tracks was manually counted at each specified distance. For Fig. 1b, the MTOC region was first defined as the region in which the majority of both directions of comets originated, and the center of that region was designated as zero.

To quantify overall MT polarity in the outgrowing PVD dendrite, for outgrowth stage, the imaging region was the whole process while the polarity quantification was done in the proximal dendrite region which is within 50μm from cell body and excluding the MTOC region; for the mature dendrite, the quantification is also done in the proximal dendrite.

To quantify the co-localization between GIP-2 and RAB-11.1in different worms (Fig. 2h), a 30 × 20 pixel ROI was drawn in the growth cone region first and then was duplicated to independent images for both GIP-2 and RAB-11.1 channels, then a random GIP-2 or RAB-11.1 localization image was generated by ImageJ JACop plugin, and then the thresholded Mander’s split colocalization coefficients(tM) values were measured by the Colocalization Threshold plugin which can generate the threshold automatically between GIP-2 and RAB-11.1, GIP-2 and randomized RAB-11.1, randomized RAB-11.1 and GIP-2.

To quantify the co-localization between GIP-2 and RAB-11.1 over time in the same worm, the thresholded Mander’s split colocalization coefficients, Pearson’s correlation coefficient (Rcoloc) and the scatterplot files were all generated by the Colocalization Threshold plugin for 100 frames in both the growth cone and cell body region.

To measure the GIP-2 cluster diameter and distance from t=0 (Fig. 3e,3f, 3g, 4g, 4h), as shown in Fig. 3e, a segmented line was draw along the GIP-2 moving track, and then the plot file for all time frames were generated in ImageJ and exported to an excel file to get the intensity value of different positions along the line and also the intensity values at different time points. Then the same segmented line was moved to a nearby region to get the background intensity value and also exported to an excel file. The final GIP-2 intensity value was the original intensity value minus the background intensity value. The pixel value of the cluster diameter at one given time point D_t_ was the distance between the first (P_first_) and last (P_last_) position in which the intensity value is above the 50% maximum intensity value in the same time point. The pixel position of the cluster P_t_ was the center of P_first_ and P_last_ (Fig. 3F). And then

Diameter (μm) =0.109* D_t_

Distance from t=0 (μm) =-0.109*(P_t_-P_0_) (0.109μm/pixel for the 100X objective calibration)

The RAB-11.1 distance from t=0 (Fig. 3n and o) was calculated the same way as GIP-2, but the threshold was set to above 70% maximum RAB-11.1 intensity, as the mCherry::RAB-11.1 showed a higher background than GIP-2.

To measure the GIP-2 cluster maximum intensity dynamics during movement (Fig. 4j), the background of the movie was subtracted using the ImageJ subtract background function, then the maximum intensity of both the green GIP-2 channel and red PVD morphology marker channel at different time points were measured using the ImageJ ROI multiple measure function. The green GIP-2 intensity was divided by the red PVD morphology channel intensity to correct for focus change or photo-bleaching, and the normalized GIP-2 maximum intensity at different time points we calculated by setting the start value as 1.

The area under the different maximum intensity curves were calculated using Prism.

Any movie that showed a slight movement was aligned by ImageJ StackReg plugin before analysis.

### *D. melanogaster* methods

The previously-generated strain *Rluv3-Gal4, UAS-EB1::GFP*^35^ was used to investigate the organization of MT minus ends in *D. melanogaster* class I da sensory neurons. Preparation of embryos was carried out as previously described^35^. Briefly, stage 15 embryos were quickly de-chorionated, washed in water, and mounted with halocarbon oil with one layer of double-sided tape as a spacer. Imaging of *D. melanogaster* was performed with a FV1200 Laser Scanning Microscope (Olympus) equipped with a 473 nm laser, and a PLAPON 60XO NA1.42 (Olympus) objective. Time-lapse images were acquired at 5% laser power at 5x zoom, imaging three z-sections spaced 0.6 µms apart, with 2.4 seconds between frames, for four minutes. Kymographs were prepared using ImageJ using an average projection of imaged z-sections.

---

**Movie 1.**

EBP-2::GFP comets reveal an MTOC in the outgrowing anterior dendrite growth cone of a wt PVD neuron in *C. elegans*.

**Movie 2.**

EB1::GFP comets reveal an MTOC near the outgrowing dendrite tip of vpda in *D. melanogaster*.

**Movie 3.**

EBP-2::GFP comets in the outgrowing PVD dendrite of a GIP-1^PVD(-)^ animal shows plus-end-out MTs

**Movie 4.**

EBP-2::GFP comets in the outgrowing PVD dendrite of an *unc-116(e2310)* mutant shows predominantly plus-end-out MTs.

**Movie 5.**

EBP-2::GFP comets in the cell body of PVD in an *unc-116* mutant during dendrite outgrowth suggests an MTOC is mislocalized to the cell body.

**Movie 6.**

GIP-2::GFP shows a stereotyped movement in the outgrowing PVD dendrite.

**Movie 7.**

GIP-2::GFP does not move in an *unc-116*(*e2310*) mutant.

**Movie 8.**

GIP-2::GFP dispersion in a *dhc-1*(*or195*) mutant.

